# High resolution discovery of chromatin interactions

**DOI:** 10.1101/376194

**Authors:** Yuchun Guo, Konstantin Krismer, Michael Closser, Hynek Wichterle, David K. Gifford

## Abstract

Chromatin interaction analysis by paired-end tag sequencing (ChIA-PET) is a method for the genome-wide *de novo* discovery of chromatin interactions. Existing computational methods typically fail to detect weak or dynamic interactions because they use a peak-calling step that ignores paired-end linkage information. We have developed a novel computational method called Chromatin Interaction Discovery (CID) to overcome this limitation with an unbiased clustering approach for interaction discovery. CID outperforms existing chromatin interaction detection methods with improved sensitivity, replicate consistency, and concordance with other chromatin interaction datasets. In addition, CID also outperforms other methods in discovering chromatin interactions from HiChIP data. We expect that the CID method will be valuable in characterizing 3D chromatin interactions and in understanding the functional consequences of disease-associated distal genetic variations.

## INTRODUCTION

Physical three-dimensional (3D) chromatin interactions between regulatory genomic elements play an important role in regulating gene expression (1, 2). For example, the creation of chromatin interactions between the promoters and locus control regions of the β-globin gene is sufficient to trigger transcriptional activation, indicating that chromatin looping causally underlies gene regulation (3).

Chromatin interaction analysis by paired-end tag sequencing (ChIA-PET) is a technology for the genome-wide *de novo* detection of chromatin interactions mediated by a specific protein factor (4). In ChIA-PET, crosslinked chromatin is sonicated and then immunoprecipitated by antibodies that bind to a protein of interest, followed by proximity ligation, and sequencing (4). The paired-end tags (PETs) are then mapped to the genome to identify the two genomic locations that interact with each other. Therefore, similar to Hi-C data (5), the ChIA-PET interactions are represented by a pair of genomic locations that interact with each other. By focusing on the chromatin interactions associated with a specific protein, ChIA-PET is capable of generating high-resolution (~100 bp) genome-wide chromatin interaction maps of functional elements (6). The ChIA-PET method has been used to detect structures defined by architectural proteins, including CTCF (6, 7) and cohesin (8, 9), detect enhancer–promoter interactions associated with RNAPII (10–12), and detect interactions involving other transcription factors (4, 13). In addition, multiple studies have applied the ChIA-PET method to link distal genetic variants to their target genes and to study the structural and functional consequences of non-coding genetic variations (6, 14).

To gain biological insight from ChIA-PET data, computational analysis pipelines and statistical models have been developed (15–19). Typically, analysis pipelines start with data pre-processing that includes linker filtering and linker removal. The resulting PETs are then mapped to the genome and duplicated PETs are removed. To detect chromatin interactions, a peak-calling step (16, 17, 19) is usually used to define peak regions enriched with reads as interaction anchors, and then groups of PETs linking two peak regions are considered as candidate interactions. Finally, the number of PETs supporting a candidate interaction is used to compute the statistical significance of the interaction.

Existing chromatin interaction methods based on peak-calling (16, 17, 19) lose information at the peak-calling step by ignoring the paired-end linkage information that is indicative of chromatin interactions. For example, for an RNAPII ChIA-PET dataset that aims to detect promoter-enhancer interactions, the RNAPII signal enrichment at certain weak or dynamic enhancers may not be strong enough to be detected as a peak by the peak-calling algorithm. Thus, interactions involving weak enhancers typically will not be detected, even though there may be a sufficient number of PETs linking these enhancers to other genomic elements in the raw data. In addition, for interactions with detected anchors, the PET count quantification may be inaccurate because some nearby PETs may fall outside of the peak region boundaries. Thus, peak-calling-based approaches limit the detection of candidate interactions and can inaccurately quantify the PET count support.

We developed a novel computational method called chromatin interaction discovery (CID) that uses an unbiased clustering approach to detect chromatin interactions to address the shortcomings of peak-calling-based methods. We show that CID can be applied to both ChIA-PET and HiChIP data and that CID outperforms existing peak-calling-based methods in terms of sensitivity, replicate consistency, and concordance with other chromatin interaction datasets.

## MATERIALS AND METHODS

### Segmentation of PETs

First, CID groups all the single-end reads that are within 5000 bp of each other into non-overlapping regions. The maximum DNA fragment size in the ChIA-PET protocol is estimated to be about 5000 bp (16). Therefore, two groups of reads that are more than 5000 bp apart are expected to belong to independent interaction anchor regions. Next, for each region, we group PETs whose left reads map to the region into groups where the right reads of the PETs map to independent anchor regions that are at least 5000 bp from each other. We then further split the PET groups if the left reads of the PETs in a group can be split into independent anchor regions. This process iterates until the PET groups cannot be further split. The result of this segmentation step is that millions of PETs are split into small non-overlapping groups that typically contain less than 10,000 PETs.

### ChIA-PET and HiChIP Datasets

ChIA-PET datasets (17 datasets associated with protein factors such as POL2RA, CTCF, and RAD21) from various cell types (9, 11, 14) (Supplementary Table S1) were downloaded from the ENCODE Project portal (https://www.encodeproject.org/). FASTQ files of both biological replicates were pre-processed and aligned to the hg19 genome using the Mango pipeline (19). The fastq and pre-processed SMC1A HiChIP data from GM12878 cells (20) were downloaded from NCBI GEO portal (GSE80820). BEDPE files from ChIA-PET and HiChIP datasets are used as inputs to CID.

### Mango and ChIA-PET2 pipelines

Mango (version 1.2.1) (19) was downloaded from https://github.com/dphansti/mango. Additionally, we installed the dependencies *R* (version 3.4.4), *bedtools* (version 2.26.0), *macs2* (version 2.1.1.20160309), and *bowtie-align* (version 1.2). Mango was executed with the default parameters and the flags *verboseoutput* and *reportallpairs* were set. For data sets that were generated with the ChIA-PET Tn5 tagmentation protocol, additional parameters recommended by the author were used: –keepempty TRUE –maxlength 1000 –shortreads FALSE.

The BEDPE files generated by Mango after step 3 were also used by the ChIA-PET2 and CID pipelines in order to examine the differences in the subsequent peak calling and interaction calling steps.

ChIA-PET2 (version 0.9.2) (17) was obtained from https://github.com/GuipengLi/ChIA-PET2. The default setting for all parameters were used, except that the starting step was set to 4 to start the analysis from Mango-derived BEDPE files.

### hichipper pipeline

The HiChIP raw fastq files were initially processed with HiC-Pro (21) (https://github.com/nservant/HiC-Pro) using default settings except specifying MboI instead of HindIII digestion. Subsequently, hichipper (22) (https://github.com/aryeelab/hichipper) was used to analyze the HiC-Pro output, specifying EACH,ALL as the peaks option and providing the MboI BED file for restriction fragments.

### Replicate consistency analysis

For each dataset we counted the number of interactions that are present in both replicates. Jaccard coefficients are then calculated by dividing the intersection of interactions in replicates 1 and 2 by the union of interactions in both replicates. Interactions in replicates 1 and 2 were considered identical, if both interaction anchors overlapped between replicates or the gap between them was less than 1000 bp.

In situations where the ranking of interactions mattered (e.g., fraction of replicated interactions in the top *n* interactions), interactions were sorted in ascending order of their false discovery rate (FDR) and posterior probability (if there were tied FDR values).

### Functional annotation of interaction calls

The GENCODE 19 gene annotation (23) was used to generate the promoter annotations. Each transcription start site (TSS) is expanded to 2.5kb up/downstream to define a promoter. We used ChIP-seq peak calls of H3K27ac histone modification, which associates with active enhancers, as the enhancer annotations. The set of broad peak calls of H3K27ac ChIP-seq data from K562 cells was downloaded from ENCODE project website (accession ENCFF931VAQ). For the interaction calls from all the methods, a call is considered annotated as an enhancer-promoter interaction if one anchor region of the interaction overlaps with a promoter annotation and the other anchor overlaps an enhancer annotation.

### Hi-C loop overlap analysis

Hi-C loop calls for GM12878 and K562 cells (24) were downloaded from NCBI GEO portal (GSE63525, combined primary and replicate samples). The HICCUPS loop calls from SMC1A HiChIP data were downloaded from Mumbach et al. (20). The overlap between Hi-C loops and ChIA-PET and HiChIP interaction calls were computed using *pairToPair* in *bedtools* with parameters “-slop 1000 -type both -is”.

### 5C interaction overlap analysis

5C interaction calls for K562 cells were downloaded from the original study (25). The overlap between 5C interactions (tested and positive) and ChIA-PET interaction calls were computed using *pairToPair* in *bedtools* with parameters “-slop 1000 -type both -is”.

## RESULTS

### Chromatin interaction discovery (CID)

CID discovers chromatin interactions using a density-based clustering method (26) to cluster proximal PETs into interactions. CID continuously resolves anchors and thus is more flexible than peak-calling-based methods that can only discover interactions between statically identified peak regions. Once CID identifies candidate interactions, it then applies the MICC statistical model (15) to compute the statistical significance of the interactions.

CID first filters out PETs that are shorter than 5000bp because they are likely to be self-ligation PETs. CID then efficiently clusters ChIA-PET data by segmenting the total set of PETs into independent groups of proximally located PETs (see MATERIALS AND METHODS). ClD then clusters each group of PETs.

A PET *i* is represented as a two-dimensional vector [*C*_*i,L*_ *C*_*i,R*_], where *C*_*i,L*_ and *C*_*i,R*_ are the genomic coordinates of the center of the left and right reads of PET *i*, respectively, *C*_*i,L*_ < *C*_*i,R*_. The distance between two PETs is quantified as the Chebyshev distance (27) calculated from the read coordinates of the PETs:

> *Distance (PET_i_, PET_j_) = max (|C_i,L_ – C_j,L_|, |C_i,R_ – C_j,R_|)*

We employ a density-based clustering method (26) that finds cluster centers that are characterized by a higher density than their neighbors and by a relatively large distance from points with higher densities. For ChIA-PET data (Figure 1A), the density of a PET is defined as the number of neighboring PETs within a certain cutoff distance to the PET. The densities of all PETs can be visualized by plotting each PET as a point (*C*_*i,L*_, *C*_*i,R*_) on a two-dimensional space (Figure 1B). A high-density group of points on the plot suggests potential chromatin interactions between two genomic regions. After computing the density values, a delta value of a PET is defined as its distance to the nearest PET that has a higher density. For the PET with the highest density, delta is defined as the largest distance between any pair of PETs (26). By requiring cluster centers have high delta values, the clustering method prevents too many points in a high-density region from being called as cluster centers (26). The cluster centers are then the PETs with both high density and high delta values, as visualized in a clustering decision graph (Figure 1C). Following the clustering method (26), PETs are ranked by the product of their density and delta values and a PET is assigned to the same cluster as its nearest neighbor of higher density (Figure 1D). After cluster assignment, singleton clusters are interpreted as noise and are not considered as candidate interactions. Because some PET clusters may be close to each other, a post-processing step merges nearby PET clusters. The PET clusters that contain at least two PETs are then proposed as candidate interactions (Figure 1E).

**Figure 1.**
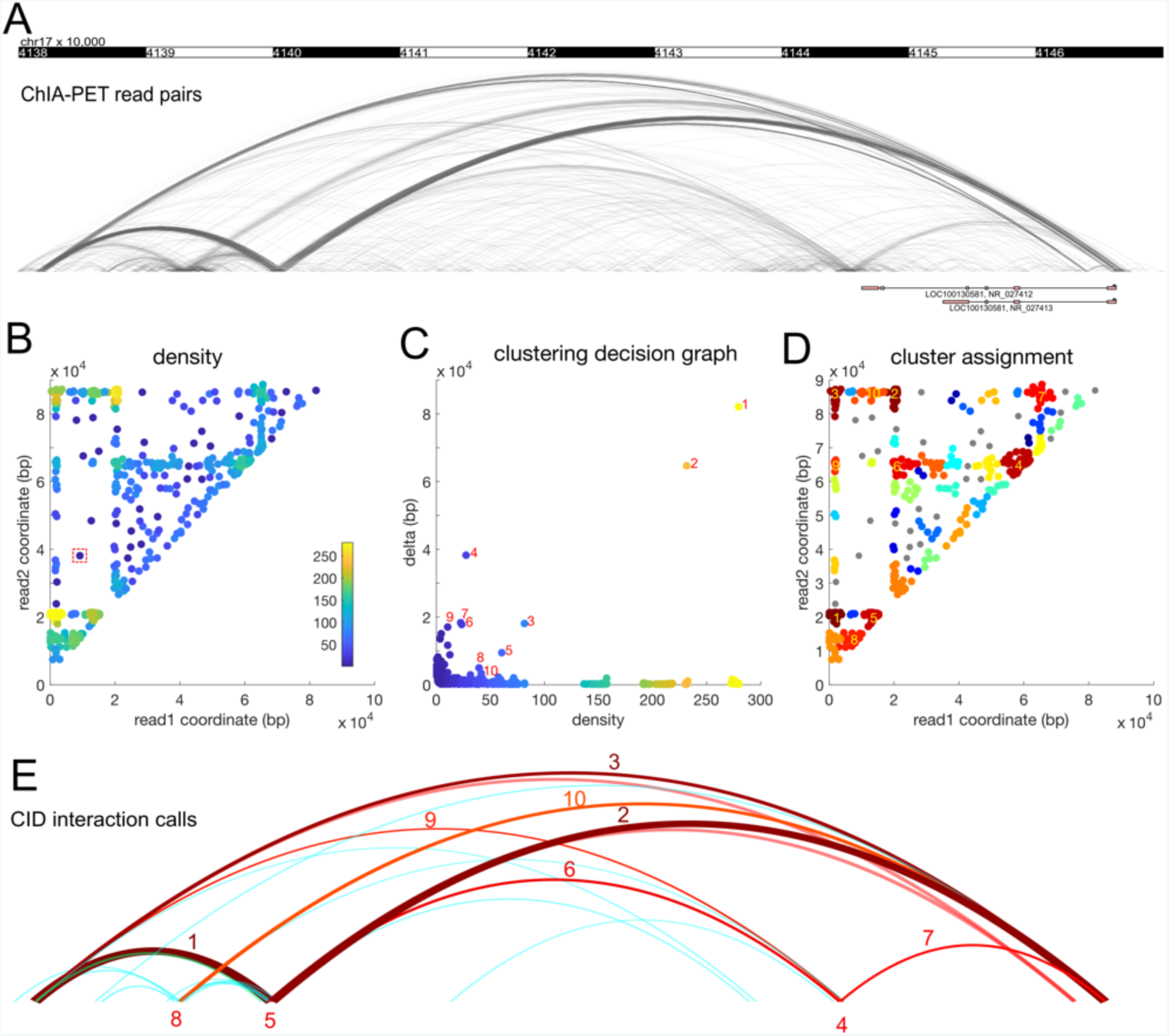
CID uses density-based clustering to discover chromatin interactions. (A) ChIA-PET interactions can be discovered as groups of dense arcs connecting two genomic regions. Each arc is a PET. (B) The PETs plotted on a two-dimensional map using the genomic coordinates of the two reads. Each point is a PET. The colors represent the density values, defined as the number of PETs in the neighborhood. The red dashed square represents the size of the neighborhood. (C) The clustering decision graph. Each point is a PET. The points with high density and high delta values are selected as cluster centers. For simplicity, only large clusters are labelled. (D) The read pairs are assigned to the nearest cluster centers. The clusters are labeled as in (C). (E) The clusters are visualized as arcs. The clusters are labeled as in (C) and (D).

CID then applies the MICC statistical model (15) to compute the statistical significance of the candidate interactions. MICC applies a Bayesian mixture model to systematically separate true interactions from random ligation and random collision noise and computes false discovery rates (FDRs) for the candidate interactions (15). The cutoff criteria for significant interactions are (1) FDR less than or equal to 0.05 and (2) PET count greater than 3. In principle, CID can the MICC, Mango, or ChiaSig (18) models to compute statistical significance of the discovered interactions. We chose the MICC model because it has been shown to be more sensitive than the Mango model (15).

### CID is more sensitive at discovering ChIA-PET interactions than peak-calling-based methods

We compared CID with two peak-calling-based ChIA-PET analysis methods, ChIA-PET2 (17) and Mango (19), and found that CID is more sensitive than these methods at chromatin interaction discovery. We tested these three methods on a widely used dataset, POL2RA ChIA-PET from K562 cells (11). We first studied the chromatin interactions called by three methods in the 400 kb genomic region downstream of the CEBPB gene. The pre-determined peak regions called by ChIA-PET2 and Mango limit the interactions that can be discovered. In contrast, CID uses an unbiased approach and discovers a substantial number of interactions that are missed by ChIA-PET2 and Mango (Figure 2A). The missed interactions are between the CEBPB promoter and non-promoter regions that have weak enrichment of reads and are not called as peaks by ChIA-PET2 or Mango. In addition, peak-calling-based methods only count the PETs that are within the peak regions and miss nearby PETs that just fall out of the peak region boundaries. In contrast, CID’s clustering approach includes all the neighboring PETs. Indeed, the PET count in the CID called interactions are higher than the same interactions called by the other methods (Figure 2A). The accurate quantification of PET counts for interactions is important for the subsequent test of their statistical significance. Many of the candidate interactions called by ChIA-PET2 and Mango contain too few PETs to reach statistical significance, yet the interactions called by CID across the same anchor regions are statistically significant because their PET counts are higher (Figure 2A). We further compared the significant interactions in the CEBPB locus between two biological replicates and found that CID called 11 replicable interactions that contain at least 9 PETs. In contrast, there are only one replicable ChIA-PET2 interaction and zero replicable Mango interactions in this region (Supplementary Figure S1). Across the whole genome, CID discovers more interactions than ChIA-PET2 and Mango (Figure 2B, Supplementary Figure S2).

**Figure 2.**
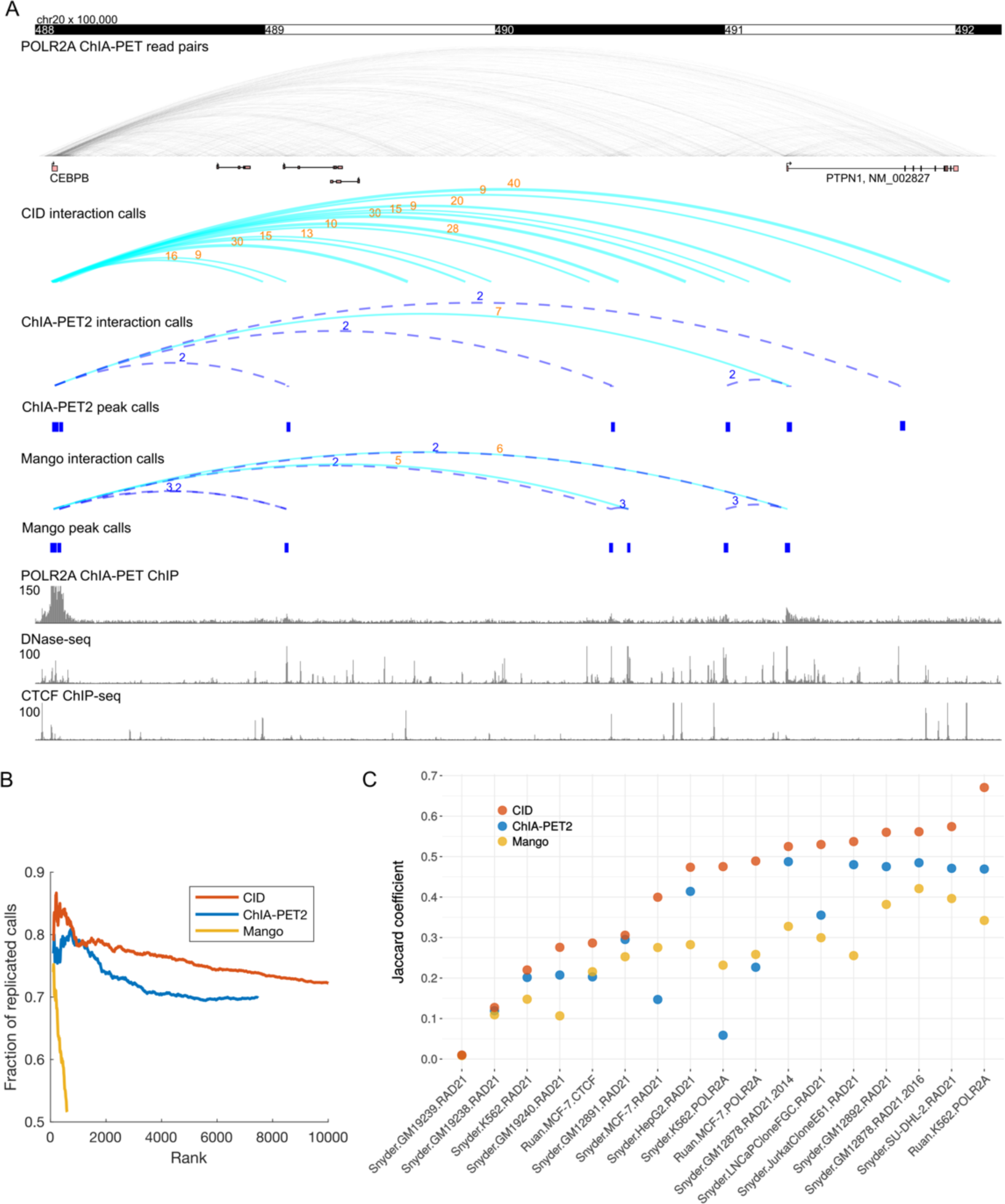
CID is more sensitive and consistent at discovering ChIA-PET interactions than peak-calling-based methods. (A) Comparison of interactions called by CID, ChIA-PET2, and Mango in the CEBPB locus using POLR2A ChIA-PET data from K562 cells. The ChIA-PET2 and Mango interaction calls are based on peak calls (shown as blue rectangles) from the same ChIA-PET data by treating PETs as single-end reads (shown as the ChIA-PET ChIP track). The PET counts of the interactions are represented as the numeric values above the arcs. For CID, only significant interactions with more than 8 PETs are shown. Dashed-line arcs represents insignificant can didate interactions. (B) Interaction calls of CID are more consistent across replicates than those of ChIA-PET2 and Mango. Accumulative fractions of replicated interaction calls are computed using top ranking interactions at increasing ranks. For CID, only top 10,000 calls are shown. (C) Interaction calls of CID are more replicable than those of ChIA-PET2 and Mango across a large set of ChIA-PET data. For each dataset, same number of top-ranking calls in replicates are used to compute Jaccard coefficient for all three methods.

Next, we investigated whether the interactions discovered by CID are functionally relevant. For the K562 POL2RA ChIA-PET data, we overlapped the interaction calls by all three methods with the annotations of enhancers (E, H3K27ac ChIP-seq peaks in K562 cells) and promoters (P, 2.5kb up/downstream of annotated TSS in GENCODE 19). An interaction is annotated as a candidate enhancer-promoter interaction if one of its anchor regions overlaps with at least one promoter or enhancer annotation and the other anchor region overlaps with at least one annotation of the opposite type (E-P or P-E). More than 80% of CID calls are annotated as candidate enhancer-promoter interactions, at the similar percentage of overlaps of calls from ChIA-PET2 and Mango (Supplementary Figure S3). Furthermore, high-ranking CID calls overlap with annotations at a higher percentage than calls from the other methods. These results suggest that CID calls reveal chromatin interactions with relevant biological function.

### CID is more consistent at discovering ChIA-PET interactions than peak-calling-based methods

In addition, CID interaction calls are more consistent across biological replicates than those of ChIA-PET2 and Mango. For each method, we compared the interactions called from biological replicates and computed the accumulated fraction of replicated calls with increasing number of top-ranking calls. For the K562 POL2RA ChIA-PET dataset, the interaction calls of CID are more replicable than those of ChIA-PET2 and Mango (Figure 2B). We further compared the replicate consistency of the three methods across a large set of replicated ChIA-PET datasets from the ENCODE project (28), which assay interactions mediated by factors such as POL2RA, CTCF, and RAD21 (a cohesin subunit), across multiple cell types. Because the numbers of interactions called by the three methods are different, for each dataset, we took the same number of top-ranking interaction calls and computed the Jaccard coefficient between the two replicates. We found that CID has higher Jaccard coefficients than ChIA-PET2 and Mango across all 17 datasets we tested (Figure 2C). Across all these ChIA-PET datasets, CID is not only more sensitive but also more consistent in discovering chromatin interactions than ChIA-PET2 and Mango (Supplementary Figure S2, Table S2). We also computed the interaction length distribution and anchor width distribution of interaction calls from Mango, ChIA-PET2, and CID for all 17 ChIA-PET data sets (Supplementary Figure S4 and S5). The interaction length distributions are similar among the tested methods. In contrast, the anchor width distribution of CID differs from other methods because CID called anchors are defined by clustered PETs instead of the peaks determined based on single-end read enrichment.

### Interactions called by CID are more concordant with HiC and 5C data than interactions called by other methods

To compare the accuracy and biological relevance of interactions detected by CID and other methods, we intersected the interaction calls with the chromatin loop calls from deeply sequenced Hi-C data (24). We first tested the concordance between RAD21 ChIA-PET interactions calls and the Hi-C loop calls in GM12878 cells. Because CID, ChIA-PET2, and Mango called different numbers of significant interactions, we focused on comparing the top 5530 interactions called by the three methods. We found that interactions called by CID overlap with more Hi-C loops than those called by ChIA-PET2 and Mango. The number of Hi-C loops overlapped with interactions called by CID, ChIA-PET2, and Mango are 2708, 1622, and 1848, respectively (Figure 3A). Similarly, for K562 cells, POL2RA interactions called by CID overlap with more Hi-C loops than those called by ChIA-PET2 and Mango. The number of Hi-C loops overlapped with the top 631 interactions called by CID, ChIA-PET2, and Mango are 88, 18, and 57, respectively (Figure 3B). In addition, the number of Hi-C loops overlapped with the top 7498 interactions called by CID and ChIA-PET2 are 396 and 83, respectively (Supplementary Figure S6).

**Figure 3.**
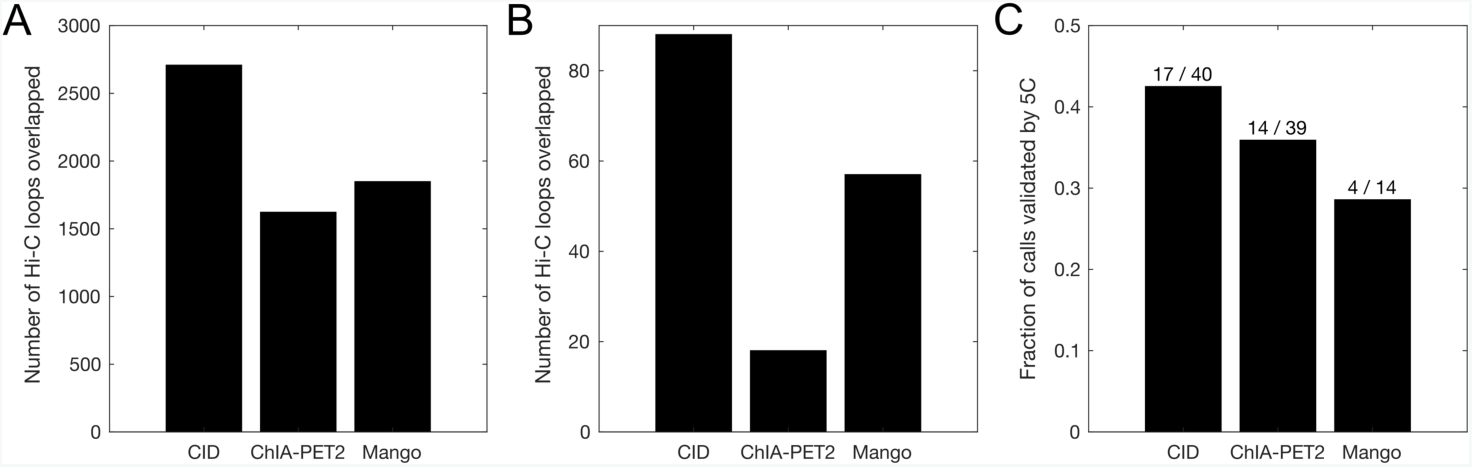
Interactions called by CID are more concordant with Hi-C and 5C data than interactions called by ChIA-PET2 and Mango. (A) Number of Hi-C loops in GM12878 cells that overlapped with top 5530 interactions called by three methods from RAD21 ChIA-PET data in GM12878 cells. (B) Number of Hi-C loops in K562 cells that overlapped with top 613 interactions called by three methods from POLR2A ChIA-PET data in K562 cells. (C) Fraction of interactions called by three methods from POLR2A ChIA-PET data in K562 cells that are validated by 5C interactions in K562 cells. The values above the bars show the number of 5C interacti ons tested positive and the number of 5C interactions tested, respectively.

We also compared the significant interactions from the three methods with 3C-Carbon Copy (5C) data mapped as part of the ENCODE project across 1% of the genome (25). For the K562 POL2RA ChIA-PET interactions called by the three methods, we compared the fraction of the interactions that are validated by 5C interactions in K562 cells. Out of 39 interactions tested by 5C that overlap the 7498 significant interactions called by ChIA-PET2, 14 were tested positive by 5C. Out of 14 interactions tested by 5C that overlap the 631 significant interactions called by Mango, 4 were tested positive by 5C. In comparison, 40 interactions tested by 5C overlap with the top 7498 significant interactions called by CID, 17 were tested positive by 5C. The fraction of positive 5C interactions are higher for CID than for ChIA-PET2 and Mango (Figure 3C).

Taken together, these results show that the interactions called by CID are more concordant with Hi-C and 5C data than interactions called by other methods, suggesting that the interactions discovered by CID are more accurate and biologically relevant.

### CID outperforms other methods in detecting chromatin interactions from HiChIP data

CID can also be applied to HiChIP (20) data for discovering chromatin interactions. HiChIP is a recently introduced method that is similar to ChIA-PET. It is an attractive alternative to ChIA-PET because it requires substantially fewer cells and a simpler protocol (20). We applied CID to a cohesin-associated HiChIP dataset (20) and found that the interactions discovered by CID are similar to a cohesin-associated ChIA-PET dataset (14) in terms of replicate consistency (Supplementary Figure S7). We then compared the results with those from hichipper (22), a peak-calling-based method for analyzing HiChIP data. We found that the interaction calls of CID are more consistent across two replicates than those of hichipper (Figure 4A). In addition, we overlapped CID, hichipper, and HICCUPS calls (20) from the same SMC1A HiChIP data with the Hi-C loops from GM12878 cells (24). Because HICCUPS only called 10255 significant interactions from the HiChIP data, we focused on comparing the top 10255 interactions called by the three methods. We found that interactions called by CID overlap with slightly more Hi-C loops than those called by HICCUPS, and significantly more than those called by hichpper. The number of Hi-C loops overlapped with interactions called by CID, HICCUPS, and hichipper are 3331, 3137, and 1507, respectively (Figure 4B). We note that HICCUPS is the same software that called the loops from Hi-C data (24). These results show that CID can also be used to detect chromatin interactions from HiChIP data.

**Figure 4.**
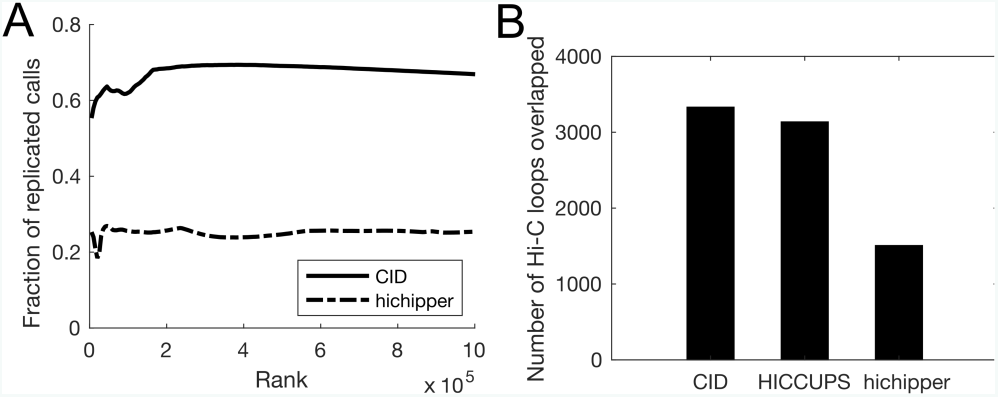
CID outperforms other methods in detecting chromatin interactions from HiChIP data. (A) Interaction calls of CID are more consistent across replicates than those of hichipper. Accumulative fractions of replicated interaction calls are computed using top ranking interactions at increasing ranks. Top 100,000 calls are shown. (B) Number of Hi-C loops in GM12878 cells that overlapped with top 10255 interactions called by CID, HICCUPS, and hichipper from SMC1A HiChIP data in GM12878 cells.

## DISCUSSION

We have demonstrated that CID is more sensitive in discovering chromatin interactions from ChIA-PET data than existing peak-calling-based methods. In addition, the interactions discovered by CID are more consistent across biological replicates and more concordant with other types of chromatin interaction data than those discovered by existing methods. We anticipate the improved accuracy and reliability of CID will be important for elucidating the mechanisms of 3D genome folding and long-range gene regulation.

We have also shown that CID can be used to detect chromatin interactions from HiChIP data. A recent study (22) showed that correction of the cut site bias of the restriction enzyme improves the detection of interaction anchors. Future development of CID with HiChIP-specific modeling of the cut site bias may further improve the detection of interactions from HiChIP data.

Cell-type-specific gene expression is often regulated by distal enhancers, and these enhancers are often enriched with disease-associated variants (29, 30). However, linking disease-associated non-coding variants to their affected genes in disease relevant tissues has been challenging due to the scarcity of long-range interaction data. With large scale on-going efforts such as the ENCODE project (28) and the 4D Nucleome project (31), high resolution chromatin interaction mapping from a wider range of tissues and cells will become available in the near future. We expect that the CID method will be valuable in characterizing 3D chromatin interactions and in understanding the functional consequences of disease-associated distal genetic variations (32).

## AUTHOR CONTRIBUTIONS

Y.G. conceived the project. Y.G. designed and implemented the CID method. Y.G., K.K., and M.C. analyzed the data. H.W. and D.K.G. supervised the project. Y.G. and K.K. wrote the paper. D.K.G. edited the paper.

## ACKNOWLEDGMENTS

We thank members of the Gifford lab for suggestions. We thank Douglas Phanstiel for suggestions on running the Mango pipeline. We thank Guipeng Li for suggestions on running the ChIA-PET2 pipeline.

## FUNDING

This work was supported by National Institutes of Health (grant 1R01HG008363 to D.K.G. and 1R01NS078097 to H.W. and D.K.G.).

